# Mutation rate variability as a driving force in adaptive evolution

**DOI:** 10.1101/354712

**Authors:** Dalit Engelhardt, Eugene I. Shakhnovich

**Affiliations:** Department of Chemistry and Chemical Biology, Harvard University, Cambridge, MA 02138

## Abstract

Mutation rate is a key determinant of the pace as well as outcome of evolution, and variability in this rate has been shown in different scenarios to play a key role in evolutionary adaptation and resistance evolution under stress. Here we investigate the dynamics of resistance fixation in a bacterial population with variable mutation rates and show that evolutionary outcomes are most sensitive to mutation rate variations when the population is subject to environmental and demographic conditions that suppress the evolutionary advantage of high-fitness subpopulations. By directly mapping a molecular-level biophysical fitness function to the system-level dynamics of the population we show that both low and very high, but not intermediate, levels of stress result in a disproportionate effect of hypermutation on resistance fixation and that traditional definitions of the selection coefficient are insufficient to account for this effect. We demonstrate how this behavior is directly tied to the extent of genetic hitchhiking in the system, the propagation of high-mutation rate cells through association with high-fitness mutations. Our results indicate a substantial role for mutation rate flexibility in the evolution of antibiotic resistance under conditions that present a weak advantage over wildtype to resistant cells.

---

The ability to predict the possible trajectories of a naturally evolving complex living system is key to describing and anticipating varied ecological and biomedical phenomena. Such predictability rests on an understanding of the *potential for evolutionary adaptability* of a given system. In asexual populations a major mechanism responsible for evolutionary adaptation under environmental stress is the generation via genetic mutations of phenotypes able to better withstand and thrive under the stressor: resistant populations arising from within a wildtype population that may “rescue” the population from the source of stress by eventually coming to dominate the population. The rate at which such resistant mutations occur and the balance between these and more deleterious mutations are major determinants of whether the population may survive and adapt to selective evolutionary pressure [1–5], an environmental stressor that targets strain variants, or phenotypes, non-uniformly. Although the baseline mutation rate in bacteria is quite low, at about ~ 10^−3^ per genome per generation [6, 7], high prevalences of mutator strains in natural bacterial populations and clinical isolates have been observed in various studies (see [8–11] for early work and [12] for a survey), and in certain cases “hypermutability”, an increase in the mutation rate over the baseline rate, was shown to result in fitness increases and faster adaptation [5, 13–18] and even be essential for survival under stress [19] by enabling genetic hitchhiking on beneficial mutations [5, 20–22]. Mutation rates can increase under environmental stress [23–26], and, in particular, hypermutability may play a significant role in the rise of antibiotic resistance [27–32].

The potential for adaptability via genetic mutations is dependent on the interplay between the ensemble of phenotypes that the system can access via mutations and the rate at which such transitions may occur within this ensemble. Phenotypes are typically characterized by some *intrinsic* measure of evolutionary fitness, such as their growth rate or lag phase, that contributes to evolutionary success, with *extrinsic* conditions, such as the probability of acquisition of this trait, initial population distribution, or resource availability, held fixed. Yet evolutionary advantage is determined by an interplay of these intrinsic and extrinsic factors, and separating these dependences while considering only a subset of them is of limited utility in establishing a global picture of a system’s evolvability potential as well as specific response to selective pressure. Here, we address both with a view to investigating the extent to which mutation rate variability drives adaptation under selective pressure.

We will assume deterministic evolution under limited resources, so that the system has a well-defined stationary state following resource saturation. We will assume in this work that the population is initially wildtype-dominated (but not necessarily exclusively so). Since at non-negligible levels of selective pressure phenotypes whose resistance to the pressure is weaker than wildtype will have very low growth, we will not keep track of such low-growth populations explicitly, but they are implicitly accounted for in our model as the loss of cells from higher-growth populations via deleterious mutations. For simplicity, we assume that such loss occurs with uniform probability *P_del_*, so that when the overall genetic mutation rate of a cell is *µ*, the rate of deleterious mutations is given by *µP_del_*. Note that when *P_del_* is high, increases in *µ* carry a higher penalty, implying that for hypermutation to be beneficial and counteract this penalty resistant phenotypes would have to be significantly advantageous either by having a much higher growth rate (intrinsic advantage) or, e.g., by occurring with a high probability or being initially present in relatively high proportions (extrinsic advantage). Here, rather than consider how variations in this advantage affect the dynamics at constant *µ* we consider the extent to which variability in *µ* affects evolutionary success at fixed levels of advantage. We quantify this effect as the stationary-state proportion of resistant mutant cells *x_r_/x_tot_* when elevated mutation rates can emerge in the population relative to when mutation rates are always kept at the low baseline *µ_bl_*:

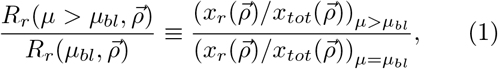

where 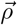 describes all parameters (initial intrinsic and extrinsic conditions) that determine evolutionary advantage in the system, i.e. that can aid the fixation of resistant phenotypes, where we define a “resistant” phenotype through its intrinsic advantage, i.e. as any phenotype exhibiting growth higher than wildtype, *g_r_* > *g_wt_* under the present level of selective pressure, a definition upon which we will expand below in the case of antibiotic resistance evolution. Importantly, since we consider here deterministic dynamics, the ratio 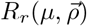 directly projects 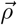 to the stationary state and should therefore be viewed as both a final outcome (at stationary-state) as well as an indicator of the evolutionary advantage conferred by the system’s initial intrinsic and extrinsinc conditions.

In our analysis we assume that phenotypes come in two mutation-rate varieties: baseline *µ_bl_* and some elevated mutation rate *f* × *µ_bl_*, where *f* > 1, whose range will be explored below. To gain conceptual insights into how general features of the phenotype ensemble impact the role of hypermutation in the system we ignore the possible effects of clonal interference and make the simplifying assumption that the resistant spectrum of the phenotype ensemble is dominated by a single growth rate *g_r_*. The resulting system and the allowed single-step transitions between the four subpopulations is shown in Fig. 1, where we note that the resistant population sizes may be set to zero in the initial state. Growth-altering mutations occur, similarly to deleterious mutations as described above, with a rate per generation that is a product of their probability of spontaneous mutation *p_wt,r_* and overall mutation rate. Since only one resistant phenotype is considered, backward and forward mutations are assumed to occur at the same rate. Mutation rate-altering mutations are assumed to occur with uniform rate *r_µ_*, and since we assume that *r_µ_* > 0 independently of any selective pressure, logic should dictate that some proportion of the initial-state population already exhibits elevated mutation rates. We assume in all that follows that cells with elevated mutation rates constitute 1% of the total initial population (distributed in proportion to the phenotype distribution) and a corresponding rate at which hypermutation-conferring mutations occur of 0.25% of cells per generation (see the Supplementary Information for an extended discussion of these parameter choices). Under the assumption of deterministic evolution in a resource-limited environment the time evolution of the population level *x_k, α_* of a phenotype with growth rate *g_k_* and mutation rate *f*_*α*_*µ*_*bl*_ is given by

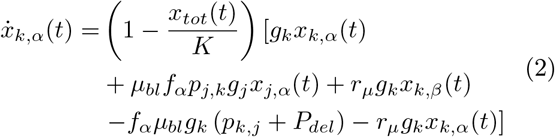

where *j ≠ k, j, k* ∈ {*wt*, *r*}, *α* ≠ *β*, and a stationary population distribution is established when the total population size *x_tot_* = Σ_*m*, *γ*_ *x*_*m*, *γ*_ reaches the resource capacity *K*. Note that faster growing phenotypes will also produce exponentially more deleterious mutants as a result of their more frequent divisions, resulting in the previously noted fitness tradeoff. The four-dimensional system (Fig. 1) of Eqn. (2) is given explicitly in the Supplementary Information.

**FIG. 1:**
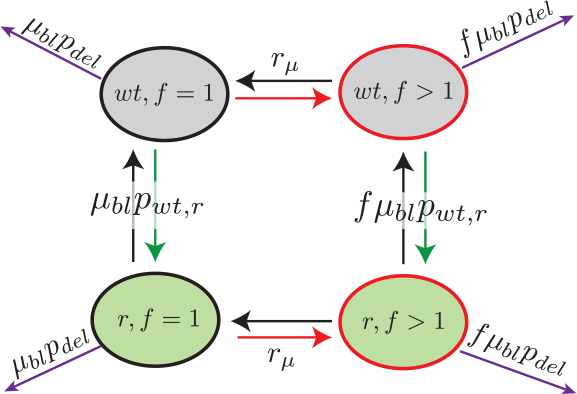
Schematic indicating the allowed single-step transitions and their rates between phenotypes.

By varying different evolutionary advantage-determining components of 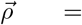 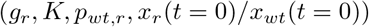 we show that for a fixed elevated mutation rate (*f* > 1) the level of advantage [^1^ We note that while the resource capacity appears a *priori* to be a non-selective environmental stressor (given the assumption made here that resource utilization among phenotypes is uniform), due to the exponential growth phase involved in the evolution of the system (2), higher resource capacity puts off the time of resource saturation, thus compounding the advantage enjoyed by phenotypes with higher growth rate *g_k_*.] due to any *ρ_i_* is negatively correlated with the positive impact of hypermutation: as *R_r_*(*ρ_i_*, *f* = 1) decreases, the extent of resistance fixation owing to hypermutation, *R_r_* (*ρ_i_*, *f* > 1)/*R_r_* (*ρ_i_*, *f* = 1), increases. The deviation from parity grows exponentially as the level of advantage decreases and becomes effectively negligible at high advantage (*R_r_*(*ρ*_*i*_, *f* = 1)→ 1). This is shown for a particular choice of *f* = 150 in Fig. 2a. By averaging over individual-p_i_ interpolations (Fig. 2b) and varying *f* (Fig. 2c and 2d) we observe that the largest impact of the presence of elevated mutation rates in the population is under initial condition combinations that, due to any one or multiple advantage-determining parameters, result in a resistant phenotype ensemble dominated by low-advantage mutants. In these circumstances the evolutionary advantage of the resistant cells may be insufficient to establish these populations in high proportions due to competition for limited resources, and certain increases in the mutation rate may thus be critical for adaptation, even at the cost of increased deleterious mutations. When initial conditions lead high-advantage resistant mutants to dominate the ensemble, mutation rate increases offer negligible to negative benefit. The high growth rate of these populations and hence frequent cell divisions imply that increases in their mutation rate also drive approximately-exponentially increases in deleterious mutations, and that when a strong advantage exists the baseline-mutation phenotype will thus rise to fixation faster than its hypermutant counterpart. For any level of advantage, there exists an optimal mutation rate yielding the highest proportion of resistant d (green curves in Fig. 2c and 2d). Increasing the mutation rate up to this rate provides substantial benefit for lower-advantage mutants, and further increases lead to diminishing (albeit more gradually) returns due to the tradeoff with an increased loss caused by deleterious mutations. We see (Fig. 2c compared to 2d) that the level of evolutionary advantage past which there is no gain from hypermutation is fairly robust to variations in the rate of deleterious mutations (*fµ*_*bl*_ *P*_*del*_), but a lower *P_del_* extends the range of mutation rates conferring benefit, as in that case there is little loss to deleterious mutations even at high *f*.

**FIG. 2:**
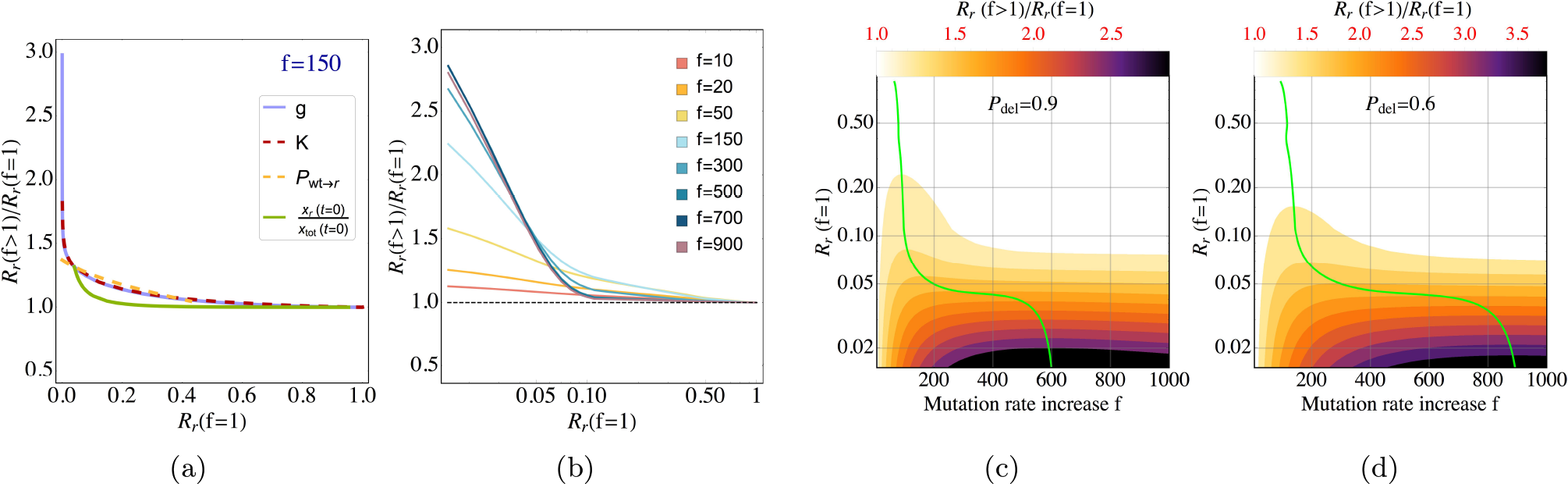
The positive effect of hypermutation (*R_r_*(*f* > 1)/*R_r_*(*f* = 1)) is negatively correlated with the baseline-mutation rate evolutionary advantage *R_r_* (*f* = 1) of the resistant mutant and is optimized at a mutation rate increase (*f*) that depends on the extent of advantage, diminishing upon further increases in this rate. (a) *R_r_* (*f* > 1)/*R_r_* (*f* = 1) versus *R_r_* (*f* = 1) curves for individual advantage-determining parameters 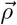 at *f* = 150. (b): averages of the single-ρ_i_ curves in the appropriate ranges put through a low pass filter for smoothness shown at multiple values of *f*. The dashed black line (parity in *R_r_* at baseline and elevated *f*) indicates the point of no benefit from hypermutation; initial increases in *f* yield significant benefit at low-advantage conditions, which decreases and eventually becomes negligible at high-advantage conditions (*R_r_* (*f* = 1) → 1). At very high *f* (here *f* ≳ 700) even low-advantage mutants experience diminishing benefit from hypermutation. (c) and (d): *R_r_* (*f* > 1)/*R_r_* (*f* = 1) contours corresponding to plot (b) in *f* — *R_r_* (*f* = 1) space. The green curve shows the optimal (i.e. yielding highest *R_r_* (*f* > 1)/*R_r_* (*f* = 1)) mutation rate increase factor f as a function of *R_r_* (*f* = 1). The probability of deleterious mutations was set at *P_del_* = 0.9 for plots (a)-(c) and at *P_del_* =0.6 in plot (d). When held constant, 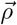 parameters were set at *g_r_/g_wt_* = 3, *K/x_tot_*(*t* = 0) = 10^2^,*p_wt, r_* = 0.01, and *x_r_* (*t* = 0) = 0. Wildtype E. coli growth was set at *g_wt_* = 0.34 h^−1^ and *µ_bl_* = 2 × 10^−10^ × *N_g_*, with *N_g_* the size of the E. coli genome.

The main force driving evolutionary adaptation, the selective pressure to which a system is subjected, typically modifies the intrinsic evolutionary advantage of phenotypes. Here, we focus on antibiotic growth inhibitors as the source of selective pressure. Motivated by work [33] on the response of E. coli to variations in the dosage of trimethoprim, a competitive inhibitor of dihydrofolate reductase, we assume a hyperbolic decay functional dependence for the growth rate *g* on the inhibitor concentration [*I*]

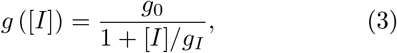

where *g*_0_ is the growth rate in the absence of an inhibitor and *g*_I_ controls the extent to which the population may grow in the presence of the inhibitor. In [33] this functional dependence, with *g*_0_ and *g_I_* given explicitly as functions of various protein biophysical and cellular properties, was shown to agree with experimental measurements for several mutant phenotypes over a range of [*I*], and similar methods can in principle be used to derive *g*_0_ and *g_I_* from biophysical principles for a wider range of biologically-relevant scenarios.

We utilize Eqn. (3) to quantify the dependence on the selective pressure [*I*] of (i) the extent to which hypermutation drives resistance fixation by computing the ratio of Eqn. (1) as a function of [*I*] with the *g_r_* parameter of 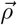 now substituted by fixed *g_r_*, _0_ and *g_r, I_* and (ii) the extent to which resistance drives hypermutation fixation via genetic hitchhiking on resistant cells by computing the stationary-state proportion of hypermutant cells of any phenotype in the presence of [*I*] (i.e. that confers positive evolutionary advantage to resistance cells) relative to in its absence:

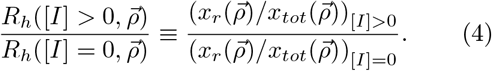

Figs. 3a and 3b show, respectively, contours of *R_r_* (*f* > 1) / *R_r_* (*f* = 1) and *R_h_* ([*I*] > 0) / *R_h_* ([*I*] = 0) in a two-dimensional space of *f* and [*I*]. We see that at low levels of inhibition, where *g_I_* carries little weight and mutant and wildtype growth rates *g*([*I*]) are similar, there is substantial benefit to be gained from hypermutation. As inhibition is increased the difference between mutant and wildtype growth increases, resulting in the resistant mutant easily increasing in proportions without much benefit from hypermutation; but at yet higher levels of inhibition the role of elevated mutation rates in determining adaptation once again becomes significant. The behavior of the (intrinsic) selection coefficient *g_r_/g_wt_* − 1 is not revealing in this respect: it monotonically approaches a constant value at high [*I*]. However, the difference between *g_r_* and *g_wt_* peaks at an intermediate value of [*I*] and decreases at lower and higher values of [*I*] (Fig. 4). While the peak does not numerically coincide with the [*I*] concentrations yielding the lowest *R_r_* (*f* > 1) / *R_r_* (*f* = 1), we note that additional parameters in 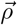 also affect this ratio. Our findings above suggest that differences in fitness *g_r_* − *g_wt_* may be a more telling representative of evolutionary advantage than the ratio *g_r_/g_wt_*, as is traditionally used to define the selection coefficient, when explicit dependence on the selective pressure is considered.

**FIG. 3:**
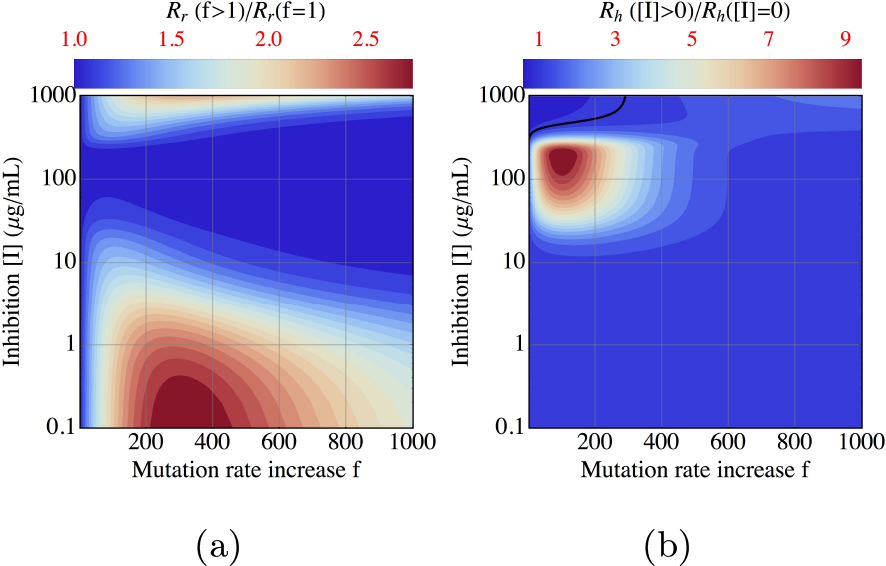
(a): Hypermutation has strong impact on resistant fixation at low and at very high levels of inhibition that is optimized at a mutation rate that depends on the inhibition level. (b): genetic hitchhiking on resistant mutations is most pronounced in intermediate levels of inhibition. The black contour (= 1) indicates no hitchhiking on resistant mutations. Parameters were set at *g*_0_,*_r_* = *g*_0_,*_wt_* = 0.34 h^−1^, *g_r,I_* = 5*g_wt,I_* where *g_wt,I_* = 3.6 µg/mL, *P_del_* = 0.9; and *K*, *p_wt,r_*, and *x_r_* (*t* = 0) as in Fig. 2.

**FIG. 4:**
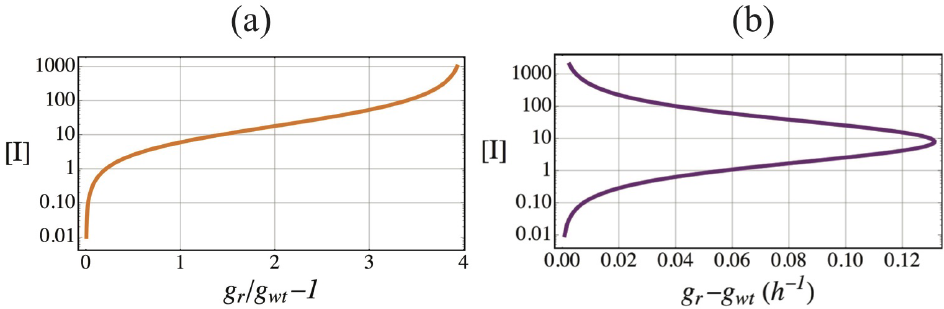
While the instrinsic selection coefficient *g_r_/g_wt_* − 1 monotonically increases as a function of inhibition (a), the difference in growth rates *g_r_* − *g_wt_* is maximized at intermediate levels of inhibition (b).

We find (Fig. 3b) that genetic hitchhiking on resistant mutations as measured by *R_h_* is most pronounced in a *f* − [*I*] phase space that up to intermediate mutation rate increases is approximately complementary to that in which hypermutation has the most pronounced beneficial effects. This effect can be explained by noting that at low inhibition, where the resistant mutant does not have significant advantage over wildtype, the acquisition of such mutations does not drastically increase the growth rate of hypermutant cells; on the other hand, when resistant mutations are highly advantageous (high inhibition), the baseline-mutation resistant mutant rises to fixation largely unaided by hypermutation, which under finite resources limits the growth potential of other subpopulations (resistant hypermutants). We note that the range of mutation rates at which we observe hitchhiking to be strongest is in keeping with experimental observations (see [12] for a review and [34] for additional recent data) of a 𝒪 (10^1^ − 10^2^) increase over baseline in E. coli clinical isolates (with some data pointing to a nearly 𝒪 (10^3^) in certain cases [35]).

In obtaining the results presented here we assumed deterministic dynamics. While mutations typically arise randomly and can introduce a large degree of stochasticity into the dynamics, deterministic evolution can provide important insights into processes with varying degrees of stochasticity: large populations are expected to sample a large extent of the available mutational phase space (with infinite populations sampling every possible configuration, or genotype), and experimental work [36] on evolutionary pathways in E. coli to drug resistance found similar mutational trajectories across populations evolved in parallel. Our deterministic results, moreover, suggest that stochastic fluctuations in the mutation rate can have an outsized effect on the stationary state of the system under a broad range of conditions that suppress the evolutionary advantage of emergent resistant populations. Knowledge of the effects of these conditions in conjunction with a quantitative understanding of how changes in a controllable selective pressure, such as we modeled here in the case of a growth inhibitor, are crucial for forming informed predictions on how variations in this main driving force of adaptation affect the dynamics of complex, high-dimensional systems and on how to best minimize the effects of stochastic fluctuations to establish a desired evolutionary outcome, such as a clinical antibiotic protocol minimizing the risk of resistance evolution.

## Acknowledgments

We are grateful to João Rodrigues for providing data on E. coli growth curves and to Michael Manhart for helpful discussions. We acknowledge support from NIGMS of the National Institutes of Health under award numbers 1R01GM124044-01 and 5R01GM068670-14.

